# KDM5B expression is suppressed by MYC in a negative feedback loop to promote cell survival

**DOI:** 10.1101/2025.04.12.648490

**Authors:** Tilottama Chatterjee, Ethan Beffert, Gabrielle S. Dewson, Delaney Sullivan, Benedict Anchang, Rosalie C. Sears, Daniel F. Liefwalker

**Author notes:** Corresponding author: Daniel F. Liefwalker, Oregon State University. Authors contributed equally to this work (Tilottama Chatterjee, Ethan Beffert).

## Abstract

Both c-MYC and KDM5B are global regulators of gene transcription, cell identity, and master transcription factors in hematopoietic malignancies. In this study we link these two critical factors in a negative feedback loop that controls apoptosis and cell survival. We find the MYC specifically downregulates KDM5B, thereby inhibiting its transcriptional repressive functions. Loss of KDM5B abrogates the cell death response to MYC withdrawal in MYC-dependent cells, indicating KDM5B mediates cell death. We find that KDM5B regulates the MYC network, and specifically demethylates the MYC locus. In summary we have discovered a negative feedback loop between MYC and KDM5B. The MYC-dependent suppression of KDM5B leads to a global increase in H3K4me3 methylation, transcriptional activity, and increase cell survival and tumor progression.

## Background

*c-*MYC (MYC) is a Transcription Factor (TF) that heterodimerizes with MAX to bind promoter regions and activate transcription across the genome^1^.

MYC activity is implicated in over 70% of human cancers^2^, and is known to influence all of the hallmarks of cancer phenotypes^3^. MYC is intrinsically disordered lacking pockets and enzymatic active sites to target with small molecules. The multivalent functions of MYC coupled with the vast number of genes regulated by MYC have confounded therapeutic approaches.

It has long been known that MYC-dependent apoptosis programs exist^4–6^, although no specific pathway or mediator of cell death has been identified. Therefore, ancillary approaches to targeting MYC activity are of clinical importance. Current understanding of MYC-dependent reprogramming to evade cell death include a double hit of *MYC* amplification accompanied by a BCL2 or BCL6, or a MYC^T58A^ mutation preventing degradation^7^, or loss of the tumor suppressor p53^8–10^. Restoring a *non-mutated* status in cancer cells is currently unfeasible as a therapeutic approach. Therefore, identification of a credible target that promotes apoptosis in response to elevated MYC has been a long-standing endeavor in the field.

In our study we analyzed a multitude of conditional MYC expressing transgenic models through a novel computational approach and find several critical nodes of MYC-dependent coordination with the JmjC class of proteins. Our findings show that MYC suppresses the histone demethylase KDM5B (JARID1B/PLU1) a transcriptional repressor which erases H3K4me3 marks used to recruit transcriptional machinery^11–13^. The inverse relationship between MYC and KDM5B has not been previously reported. KDM5B is a key regulator of pluripotent potential, where KO mice are embryonic lethal^14^. KDM5B was initially identified through its overexpression in breast cancer^15–17^, and elevated KDM5B is in some hematologic malignancies^18–23^, an observation that has been extended to several other cancer subtypes^24–32^. In other studies KDM5B is shown to act as a tumor suppressor. For example KDM5B has been shown to suppress self-renewal and oncogenic potential in leukemia^33^, and exerts context dependent control over differentiation and self-renewal^34–41^. KDM5B loss phenocopies loss of retinoblastoma (Rb) in Rb-dependent models^42,43^. In triple negative breast cancer cells KDM5B restricts cell migration^44^ and associates with the Nucleosome Remodeling and Deacetylase (NuRD) complex^45^.

In this study, we find that MYC suppresses KDM5B as part of a critical node of regulation in MYC-dependent T-ALL. We also find that KDM5B regulates pro-apoptosis responses and the MYC network and is part of a part of a negative feedback loop that regulates the Myc locus.

## Results

### Discovery of critical nodes in MYC-dependent tumors with nested effects model

We began these studies by seeking common transcriptional signatures across several MYC dependent cancers. Gene expression data was obtained of primary transgenic models of cancer initiated by conditional expression MYC, and cell lines derived from the primary models^3,46–52^. These inducible models contain an ectopic Tet-off system expressed by tissue specific promoter sequences, coupled with a tTA promoter driving MYC expression, where the addition of doxycycline quenches expression of MYC. Our strategy was to treat the data set as a complex transcriptional landscape with nested features (nodes) that have a predominate impact on shaping the transcriptional landscape. We interrogated the data set to uncover underlying gene programs driving topical MYC-dependent program perturbations (MYC on vs off). We utilized a systems biology Nested Effects Model (NEM)^53,54^ approach. In this analysis, the NEM infers network relationships concealed within the transcriptional landscape and hierarchically clusters the underlying nodes that drive the observed phenotype.

We found relationships between disparate cancer models in the context of MYC on versus MYC off (MYC axis), as well as cancer subtype specific nodes. For example, the metastatic HCC in lung driven by MYC and Twist1 are nested within the signatures for primary MYC-driven HCC, primary HCC driven by MYC/Twist1^55^, and metastatic HCC in kidney and lymph node (Fig. 1A). All HCC models were part of the HCC network or set, similar to the lung adenocarcinoma model, which we interpreted as reasonable outcome for the model.

**Figure 1.**
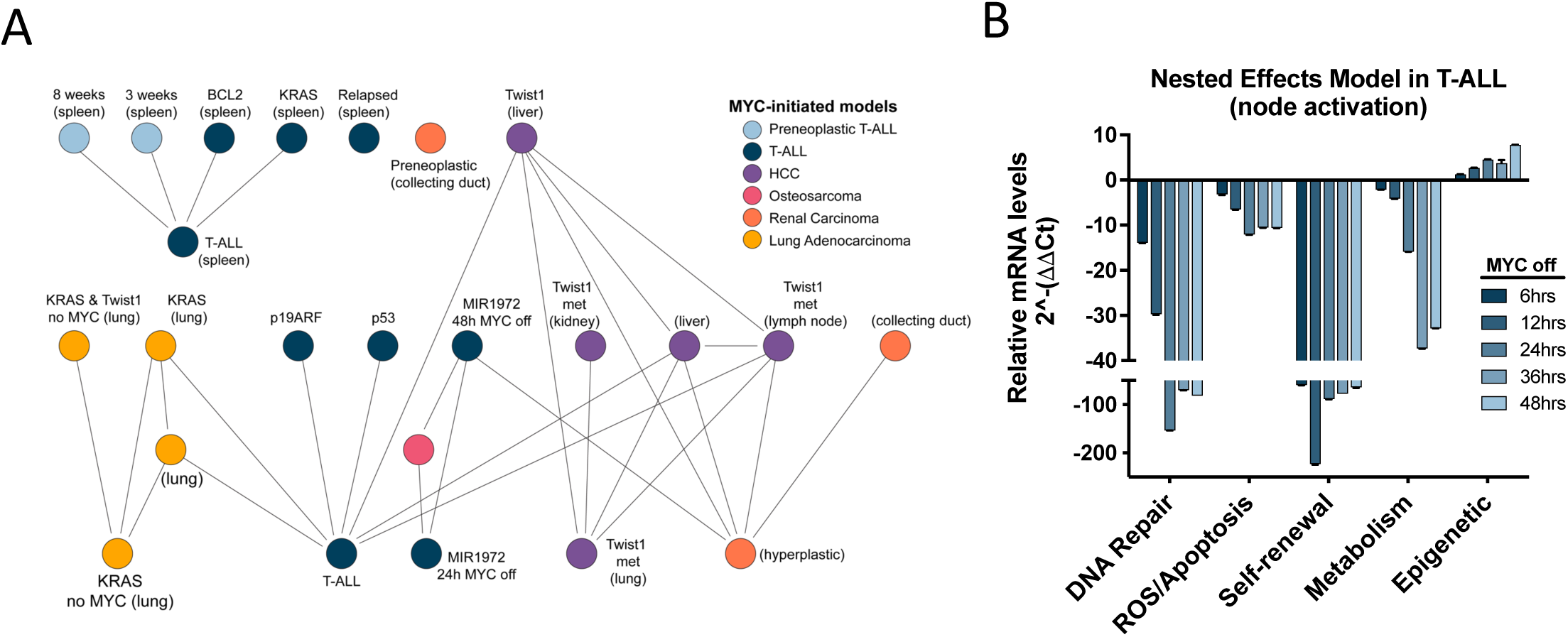
Discovery of critical nodes in MYC-dependent tumors with nested effects model. A) Transcriptional profiling of multiple conditional transgenic models of cancer were hierarchically clustered using the nested effects model. Tissue from primary models are indicated, otherwise the analysis includes cell lines from the indicated model. Lower nodes are nested within the higher nodes, and two nodes did not show nested effects. B) Applying the NEM approach to the *Eµ-tTA/Tet-O-MYC* model of T-ALL we uncovered five nodes that underly MYC-mediated programs. Expression patters of representative members of each node were examined in a MYC-off time course.

Several interesting features of specific MYC-driven cancers were also uncovered. First, the T-ALL model was nested within p53 or p19ARF null cells, a feature previously observed in MYC-dependent malignancies^6,56–59^, again confirming that our computational approach is robust. We also observed that multiple MYC-driven models converged upon the features displayed in the T-ALL model^60^. These data suggested that the hematopoietic model contains essential nodes or features that drive multiple models of MYC-driven cancers. Therefore, we applied the NEM to the T-ALL model data to discover programs that support MYC-dependent transcriptional landscapes and determined at least five critical nodes sustaining MYC-induced transformation (Fig. 1B). These five nodes include previously reported relationships (self-renewal^61^, metabolic^52^), as well as uncharacterized MYC-dependent nodes. In this study we examine the epigenetic node, which is in opposition to MYC expression.

### Oncogenic MYC suppresses KDM5B expression

A cell intrinsic model of MYC-reliant cancers is that of transcriptional amplification or dependency^1,62,63^, where all active gene programs are amplified by invasion of the transcriptome by MYC^64^. Of particular interest from the NEM results was the epigenetic node (Fig. 1B), which is in opposition to MYC expression. The suppression of this node is therefore not part of a transcriptionally amplified program or model. The implicated epigenetic node primarily contains the jumonji family of histone demethylases. In cells derived from the *Eµ-tTA/Tet-O-MYC* model, we studied JmjC cluster of histone demethylases and chromatin regulators^65^ and find that *Kdm5b* is uniquely upregulated after revoking MYC expression (Fig. 2A).

**Figure 2.**
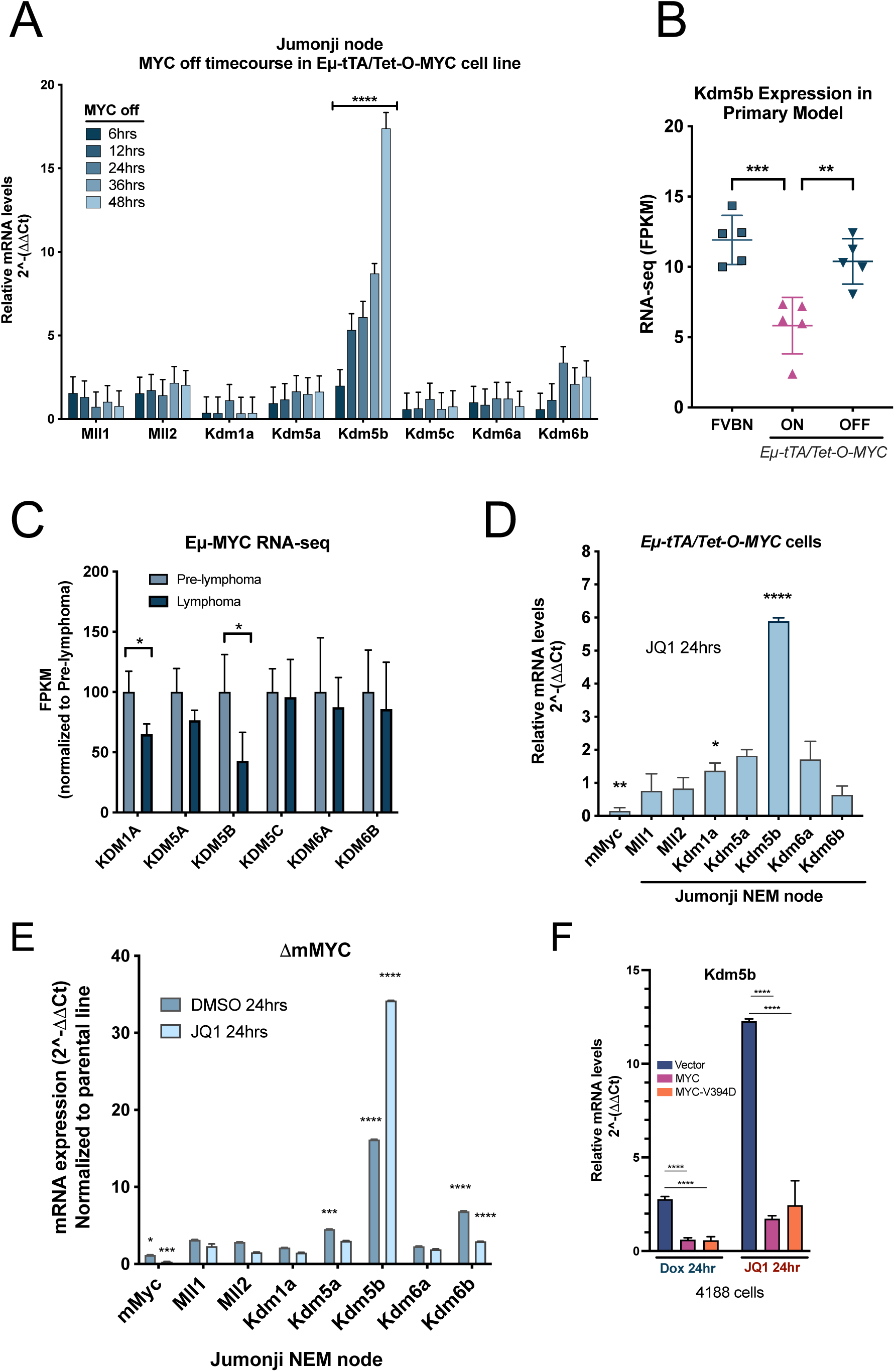
Oncogenic MYC suppresses the histone demethylase KDM5B. A) MYC-dependent suppression of JmjC cluster is specific to KDM5B. B) *Eµ-tTA/Tet-O-MYC* primary model of T-ALL RNA-seq data show reduced KDM5B levels in the MYC on condition (purple triangles) when compared to the MYC off (blue triangles) of the basal level of expression in background FVBN mice (blue squares). C) RNA-seq (GSE51011) from the *Eµ-MYC* model. KDM5B is repressed as Lymphoma develops. D) The BET inhibitor JQ1 reduces endogenous Myc levels in cell lines derived from the mouse model. KDM5B is upregulated as MYC levels decrease. E) Cas9 disruption of endogenous Myc upregulates KDM5B. Additional treatment with JQ1 potently upregulates KDM5B concomitant with enhanced reduction of Myc. F) Suppression of KDM5B is not Miz1 dependent. Cells were transfected with empty vector, MYC expression, or the Miz1 binding mutant MYC-V394D. Empty vector exhibited upregulation of KDM5B after the MYC transgene is turned off (Dox), and also upon JQ1 treatment (blue bars). When MYC is over expressed (purple bars) and constitutively suppressing KDM5B expression there is no appreciable increase in KDM5B expression. The MYC-V394D mutant does not bind to Miz1 and KDM5B expression is not upregulated (orange bars) indicating the the suppression of KDM5B is not Miz1 dependent.

KDM5B is a broad transcriptional repressor, through both associations with other complexes (NuRD)^45^ or through the demethylation of histone 3 lysine 4 reside (H3K4)^65^. The *kdm5b* gene product (KDM5B/JARID1B/PLU1) contains Arid, JmjN, JmjC, and three PHD domains. The PHD1 finger domain recognizes the H3K4 unmodified histone residue as part of the JmjC/JOR domain along with the ARID domain. The recognizes trimethylated H3K4me3 mark and exerts transcriptional repression by stripping the H3K4 methylation status to a single H3K4me1^45^, thereby acting as a transcriptional repressor.

We next examined the MYC-dependent suppression of KDM5B *in vivo*. We observe opposing expression patterns between MYC and KDM5B (Fig. 2B) in *Eµ-tTA/Tet-O-MYC* model^66^ when compared to the doxycycline (dox) treated (MYC off) and background FVBN mice. In a separate model (*Eµ-MYC* model)^64,67^ where these mice develop B-cell lymphoma we observed opposing expression patterns between MYC and KDM5B (Fig. 2C) as the malignancy progresses. Together these data demonstrate MYC suppresses KDM5B in immune intact models.

The Bromodomain Extra-Terminal (BET) containing family members bind to active enhancers and super-enhancers and contribute to cell identity and fate^68^. Inhibiting BRD4 with a small molecule (JQ1) has been shown to downregulate MYC expression^69–71^. JQ1 treatment of transgenic cells reduced endogenous MYC expression and strongly upregulated KDM5B expression (Fig. 2D). Cas9-mediated knockout of endogenous *Myc* (approximately 80%kd) was sufficient to upregulate KDM5B (Fig. 2E). Treatment with JQ1 further suppressed endogenous *Myc* and enhanced KDM5B upregulation (Fig. 2E).

Next we examined the role of Miz1, a transcriptional co-regulator of MYC-dependent programs^72,73^. In high MYC expressing cells, Miz1 is known to associate and compete for MYC binding with MAX (obligate heterodimer to MYC), and can suppress MYC target genes in this manner. To test whether Miz1 association is plays a role in repressing KDM5B we we transduced the murine 4188 cell line with MYC overexpression vectors, and a mutant MYC-V394D substitution that disrupts interaction with Miz1. We observed that the addition of both the MYC or Miz1 binding mutant continued to suppress KDM5B at similar levels even when the ectopic tet-O-MYC was suppressed (Fig. 2F), suggesting that the presence of MYC regardless of binding mutation with Miz is sufficient to suppress KDM5B. When treated with JQ1, we see that expression of KDM5B in the vector control is increased as observed in the non-transduced cells (Fig. 2D, 2E). However, with the additional constitutive expression of MYC or the Miz1 binding mutant we do not observe significant upregulation of KDM5B, suggesting that the MYC protein is suppressing KDM5B independent of the association with the transcriptional repressive function of Miz1/MYC binding. These data suggest oncogenic MYC represses KDM5B expression, either indirectly or directly.

In summary, we find that when MYC is suppressed by a transgene, JQ1 or Cas9-mediated KO, KDM5B expression is elevated suggesting that MYC negatively regulates KDM5B in murine models.

### MYC suppresses KDM5B in human hematopoietic cancers

We next examined a panel of human lymphoma and leukemia cell lines for c-MYC and KDM5B expression (Fig. 3A and 3B). We then tested the cell line panel for sensitivity to JQ1. Most cell lines were sensitive to JQ1 in a dose dependent manner, apart from the low MYC CA-46 cells (Fig. 3C). To examine the effect of JQ1 on KDM5B and MYC expression we monitored the mRNA levels and found that MYC downregulation and concomitant KDM5B upregulation was observed in JQ1 sensitive cells (Fig. 3D). These data suggest that human hematopoietic malignant cells are sensitive to disruption of MYC expression, presumably due to increased KDM5B expression.

**Figure 3.**
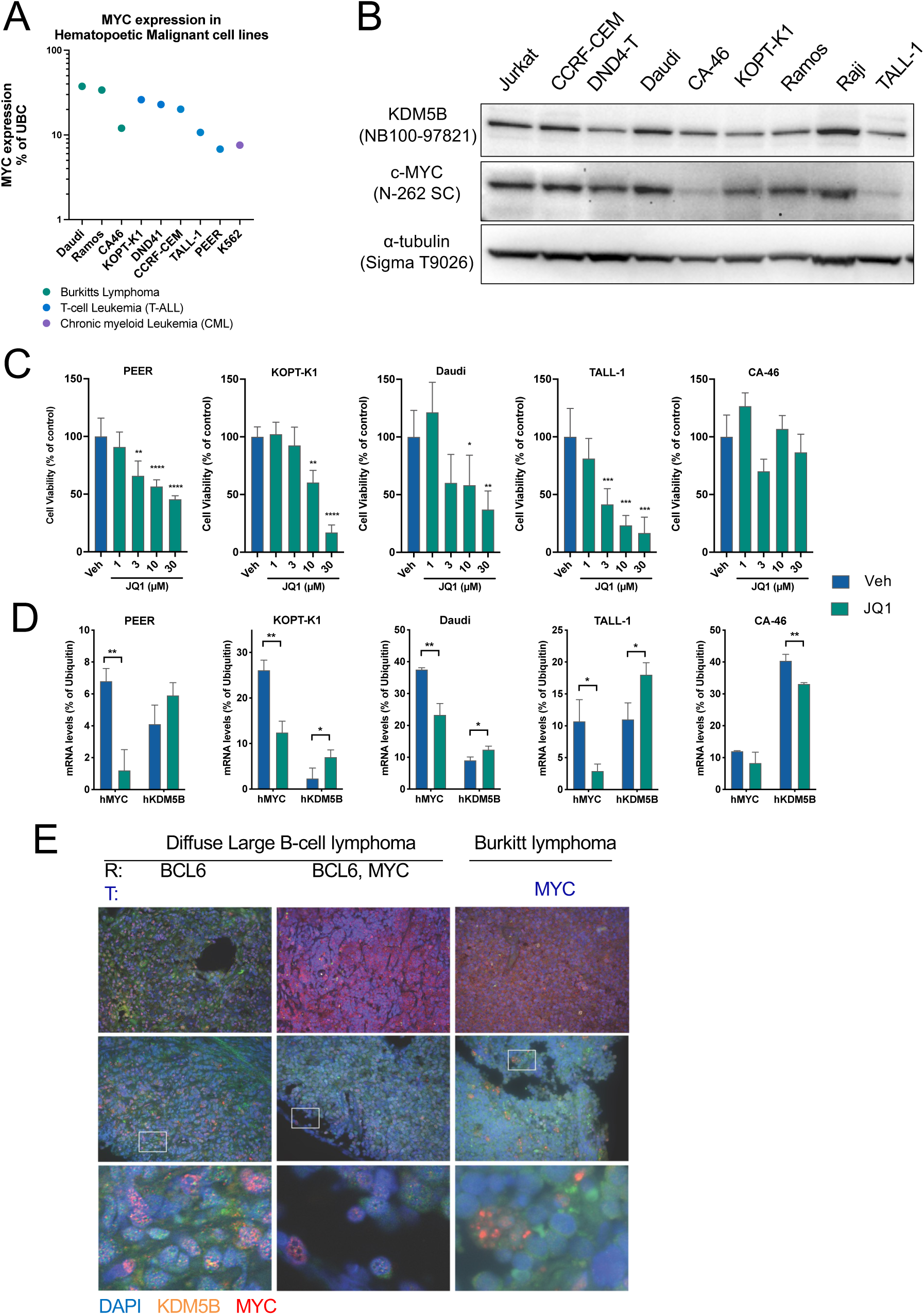
MYC suppresses KDM5B in human hematopoietic cancers. A) Transcriptional profiling of MYC in various human lymphoma and leukemia cell lines. B) MYC and KDM5B protein expression in a panel of human lymphoma and leukemia cell lines. C) A panel of human hematopoietic cell lines are sensitive to JQ treatment, with exception of the low MYC expressing CA-46 cells. D) JQ1 sensitive cells exhibit reduced MYC levels and increased KDM5B. E) Four DLBCL and two Burkitt lymphoma clinical samples (blinded study). High MYC cells and tissue show reduced or absent KDM5B. Strong KDM5B signal show reduced MYC expression. Pathology report specify rearrangement via FISH (top): (R) or translocation (T). Final row is the indicated white square.

Diffuse Large B-cell Lymphoma (DLBCL) is the most common non-Hodgkin lymphoma diagnosed in the United States. Burkitt lymphoma is a subtype of DLBCL driven by a *c-MYC* translocation event most commonly to an *IgH* locus t(8:14), leading to constitutive expression^74^. Multiple hematopoietic malignant cells display aberrant MYC signaling^7,75–82^. Blockade of apoptotic responses is a central feature of MYC-dependent hematopoietic malignancies. Therefore, we examined the expression profiles *in situ* using human tissue sections of Diffuse Large B-Cell Lymphoma DLBCL and Burkitt lymphoma.

These clinical tissues were scored by a pathologist, tested for mutations in MYC, de-identified, and provided as a blinded study. The pathology reports indicate BCL2, BCL6, and MYC rearrangements and translocations (Fig 3E). In DLBCL tissues with no MYC mutation, KDM5B and MYC are co-expressed (left column), although individual cells exhibit opposing expression patterns of the MYC/KDM5B axis. In high MYC tissues (top middle, top right) KDM5B expression is nominal or absent. In tissues not displaying ubiquitous MYC upregulation (middle row) individual cells also exhibit opposing MYC/KDM5B expression (bottom row). Taken together, KDM5B is active in cells that are sensitive to BET inhibition, and KDM5B is suppressed by oncogenic MYC in human clinical DLBCL samples.

### Loss of KDM5B enhances cell survival

Despite cancer cells acquiring additional mutations, their progression remains dependent upon the initiating oncogene^50,83^, referred to as oncogene dependency. One such endpoint to MYC withdrawal is apoptosis^3^. To examine the contribution of KDM5B in the many biological responses observed when MYC expression is revoked^3^, we utilized a CRISPR-based approach. Two sets of sgRNAs were designed targeting *Kdm5b* early exons. After clonal expansion, several clones were identified with significantly decreased *Kdm5b* mRNA as well as greatly reduced protein levels (Fig. 4A).

**Figure 4.**
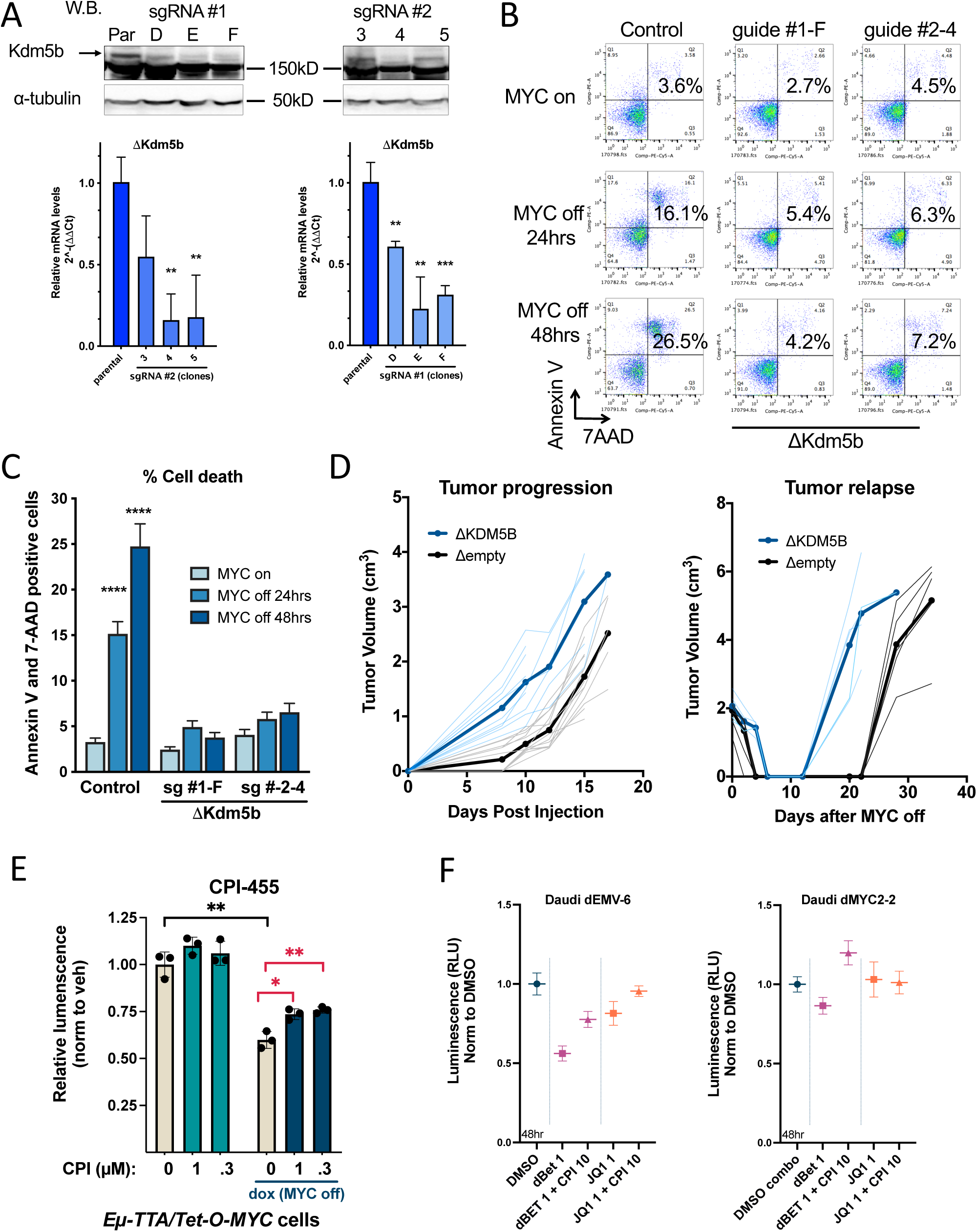
Loss of KDM5B enhances cell survival. A) KDM5B knockout *Eµ-tTA/Tet-O-MYC* cell lines. B) KDM5B KO cell lines do not undergo apoptosis when MYC expression is abrogated, in contrast to the parental control lines. C) Summary of flow cytometry replicate experiments for cell death. D) KDM5B KO lines show were subcutaneously injected into background mice and display enhanced tumor progression, with delayed collapse and escalated relapse when compared to the empty vector KO control lines. E) 4188 cells were treated with KDM inhibitor with MYC on vs off (dox). CPI treated cells show resistance to cell death when MYC expression is abrogated. F) Rescue experiments in Daudi cells show CPI partially rescues treatment with 1uM JQ1 or 1µM dBET (PROTAC). Cas9-mediated MYC KO cell line is not sensitive to BET inhibition.

Using these clones, we examined apoptosis with Annexin V and 7-AAD staining after abrogating MYC expression with doxycycline treatment (Fig. 4B). In the control cells, strong increases in apoptotic cells were observed, a phenotype arising from their dependency on MYC. The KDM5B KO cells (Fig. 4A) exhibited near complete abrogation of apoptosis after MYC withdrawal (Fig. 4B, 4C), suggesting that KDM5B plays a key role in mediating this response.

This phenotype was then interrogated *in vivo*. Syngeneic engraftment of the cell lines *in vivo* show that knockout of KDM5B confers enhanced tumor progression and a more rapid relapse in immune intact FVB/N host mice (Fig. 4D). Overall, loss of KDM5B conferred increased cell survival in MYC-dependent hematologic malignancies.

To assess the role of KDM5B enzymatic activity in regulating cell survival, we tested ^63^ CPI-455^84,85^, a selective inhibitor of histone demethylase sites on KDM5A, 5B, and 5C. Inhibition of demethylase activity with CPI-455 attenuated cell death upon MYC withdrawal (Fig. 4E), suggesting demethylase enzymatic activity has a significant role in mediating apoptosis. We expanded this rescue experiment by generating a Cas9-mediated MYC KO cell line from the Daudi Burkitt lymphoma cell line. Using both JQ1 and the PROTAC dBET, we again observed modest but significant rescue of cell populations with CPI-455 cotreatment.

These data suggest that the cell death response to collapse of MYC-dependent oncogenic programs are mediated by KDM5B, potentially through the demethylase enzymatic activity of KDM5B.

### KDM5B regulates the MYC network in opposition

As a histone demethylase KDM5B regulates the access across the genome. Multiple studies have also shown MYC exerts global influence on transcriptional programs^1^. Our research program seeks to discover the mechanistic nature of the negative feedback loop when MYC is overexpressed that leads to apoptosis^4^. Therefore we set out to discover potential opposing pathways regulated by KDM5B or MYC. We performed RNA-seq experiments on the Cas9-mediated KDM5B KO cells in the context of MYC perturbations (On versus Off) in cells derived from the *Eµ-tTA/Tet-O-MYC* model.

As expected we observed thousands of differentially regulated by KDM5B and MYC (Fig. 5A). We then performed gene ontology analysis of the genes associated with either perturbations of MYC expression or KDM5B wt versus KO (Fig. 5B). We observed significant overlap in pathways regulated by these proteins. However, nearly all of those programs were inversely correlated, suggesting a significant overlap in regulation of the mYC network. It is also interesting to note that the MYC targets were strongly correlated with both MYC and KDM5B, suggesting that KDM5B activity may also directly influence the MYC network, even when MYC expression is abrogated.

**Fig 5.**
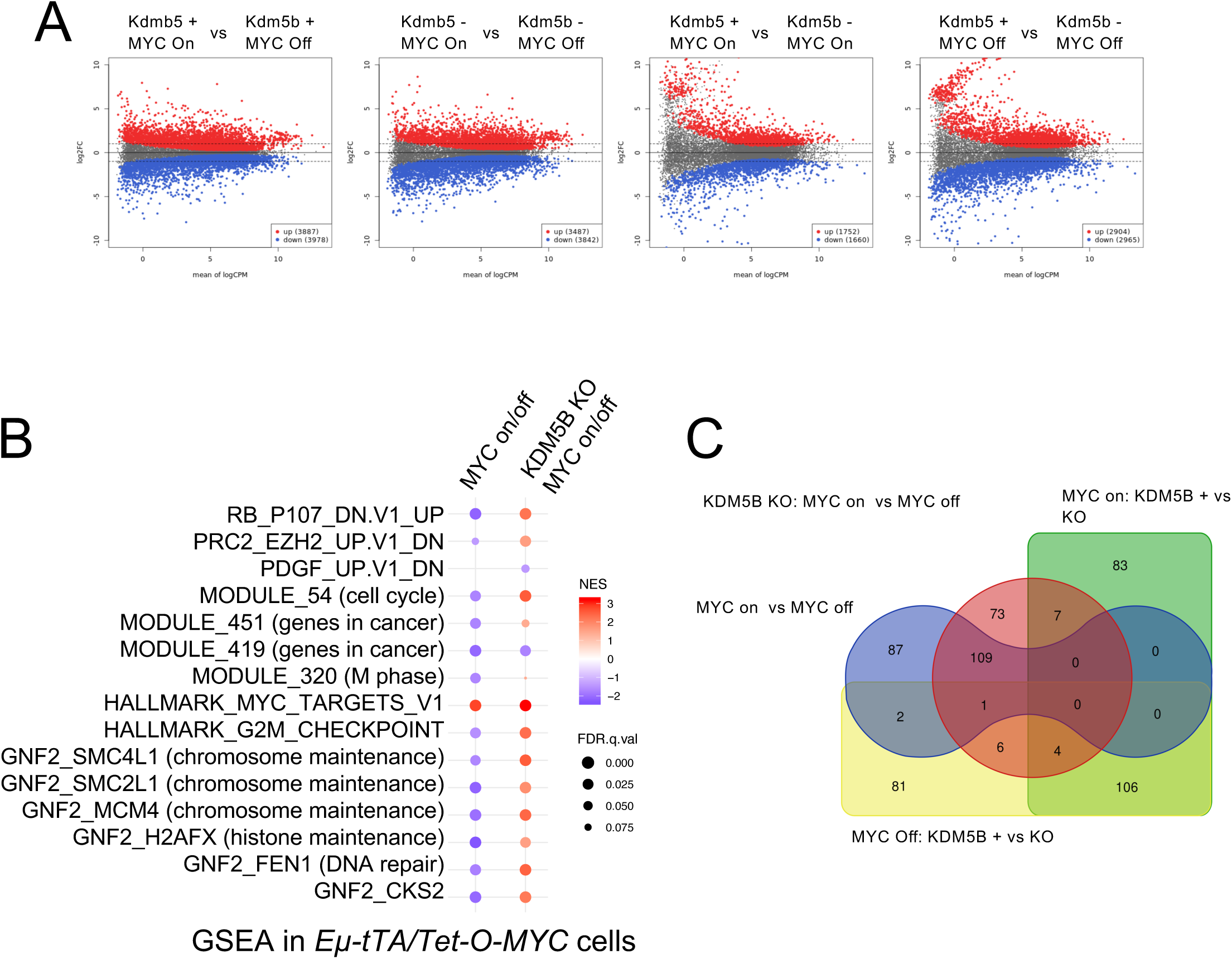
KDM5B regulates the MYC network. A) RNA-seq (MA plots) for multiple MYC/KDM5B perturbations in cells derived from the *Eµ-tTA/Tet-O-MYC* primary model reveal specific signatures and pathway dependencies. Both factors influence thousands of genes and act as global transcriptional regulators in these cells. B) Gene set enrichment analysis indicated that MYC suppresses many programs related to chromosome maintenance and cell cycle checkpoints. C) The top 200 differentially expressed genes for each perturbation (KDM5B KO vs wt, MYC on vs off) were analyzed for overlapping signatures. KDM5B KO (red) predominantly regulates MYC-dependent target genes.

To gain better understanding of specific genes that are regulated in either by both KDM5B and MYC, or by KDM5B or MYC alone we examine the transcriptional overlap of each condition. We took the top 200 differentially regulated genes from each combination and compared the gene sets (Fig. 5C). When we examine the MYC on vs off condition (blue) and compare this same condition in KDM5B KO cells (red circle) we observed 110 of the top 200 genes to be co-regulated KDM5B. Conversely, when we examined the influence of MYC in KDM5B positive versus KDM5B KO cells we observed very little overlap of MYC regulation (blue and green rectangles). This data suggests that KDM5B exerts control on the MYC network target genes.

Our RNA-seq data demonstrates, and confirms, that MYC and KDM5B are genome-wide transcriptional regulators. We were interested in the intersection of gene programs attributed to each factor. While considerable overlap exists in the gene ontology programs, the expression patterns are largely discordant. Further, our data suggest that the most prominent gene programs of MYC are also regulated by KDM5B, although MYC does not appear to regulate top KDM5B targets.

### KDM5B regulates the MYC locus, forming a negative feedback loop

While MYC is primarily characterized as a transcription factor, KDM5B exerts influence on chromatin access via histone post-translational modifications. Therefore, we were interested in examining several histone marks related to KDM5B function and transcriptional access.

Accordingly, we utilized the KDM5B KO cells to query the epigenetic landscape in ChIP-seq experiments. H3K4me3 is associated with transcriptional activation, and the demethylation of this mark is associated with transcriptional repression. Profiling of the H4K4me3 mark near transcriptional start sites (TSS) revealed a global reduction of H3K4me3 mark in the MYC off condition, likely due to the restoration of KDM5B expression (Fig. 6A) and demethylase activity. In the KDM5B KO cell lines we observed no appreciable difference in the H3K4me3 mark when comparing MYC on versus off condition. Multiple factors modify the H3K4me3 mark, which is associated with transcriptional activation. Our data data show that MYC-dependent influence on H3K4 methylation status is mediated primarily by KDM5B.

**Figure 6.**
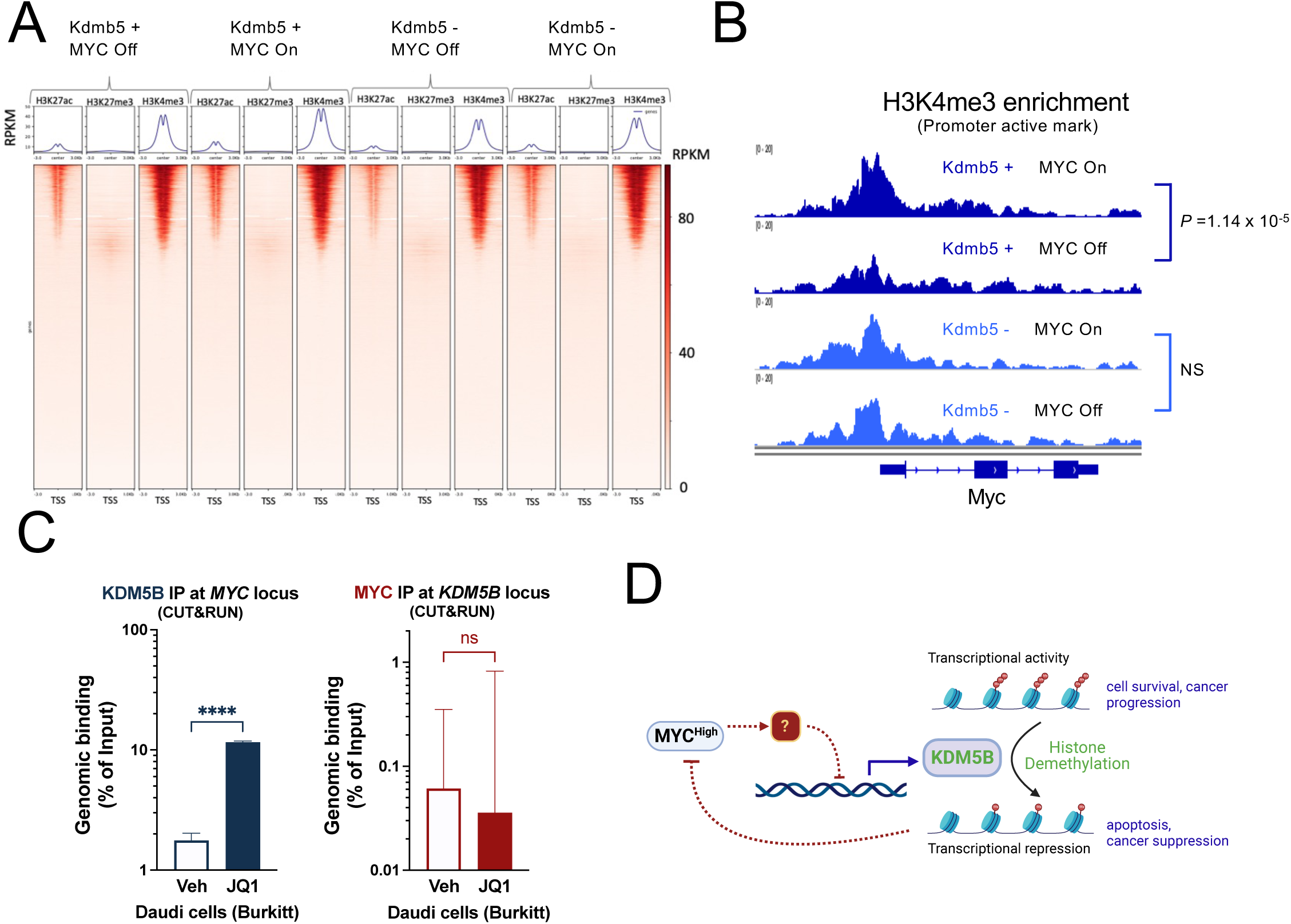
KDM5B regulates the MYC locus. A) ChIP-seq experiments were performed for H3K27ac (active enhancers), H3K27me3 (silenced chromatin/PRC2), and H3K4me3 (promoter/KDM5B activity) in each condition (KDM5B wt vs KO/MYC on vs off). Global changes in the H3K4me3 status for MYC on vs off are only observed in the KDM5B wt condition. B) ChIP-seq data reveals that the H3K4 methylation status of the *Myc* promoter region is under control of KDM5B. C) JQ1 treatment increases KDM5B binding at the *Myc* locus in the Burkitt lymphoma cell line Daudi. MYC was not significantly detected at the *KDM5B* promoter region. D) Graphical abstract. Model derived from preliminary data features mutual repression through a feedback loop, and apoptotic control by KDM5B.

We next examined the top genomic targets exhibiting statistically significant changes in H3K4me3 in the MYC on vs off conditions. Among the top hits was the MYC locus (Fig. 6B). The ChIP-seq data show a clear reduction of the H3K4me3 mark in the MYC off condition (Fig. 6B, dark blue), presumably due to the restoration of KDM5B expression and demethylase activity. Indeed, in the KDM5B null cells the differential regulation of the MYC locus is absent (Fig. 6B, light blue). Thus, the H3K4me3 mark associated gene activation is demethylated in a KDM5B dependent manner. Given that these studies demonstrate a MYC dependent suppression of KDM5B, these two factors form a negative feedback loop.

To examine this axis in human cells we again utilized the human Daudi Burkitt lymphoma cell line to directly examine the reciprocal regulation of KDM5B at the MYC locus, or MYC at the KDM5B locus (Fig. 6C). Using CUT&RUN we find robust increases in direct binding of KDM5B at the MYC locus when MYC expression is inhibited by JQ1. However, we did not detect MYC binding at the KDM5B locus in either condition, suggesting an indirect suppression of KDM5B.

Collectively, these data show that KDM5B mediates MYC-dependent influence on the epigenetic H3K4me3 mark, and that KDM5B directly regulates the MYC locus (Fig. 6D), as part of a reciprocal negative feedback loop.

## Discussion

The master transcription factor *c*-MYC plays a central role in a multitude of cellular processes. Primary roles include cell growth and differentiation, ribosome biogenesis, metabolism and transcriptional activation of the genome. MYC is one of four Yamanka factors known to reprogram cells into a pluripotent state. As an oncoprotein MYC underlies many cancer associated processes, and is implicated in up to 70% of all cancers^2^ and is implicated in all of the Hallmarks of Cancer phenotypes^3^. We set out to discover shared fundamental underlying programs that support MYC-dependent malignancies across a wide variety of MYC-dependent models of cancer (Fig. 1A). The JmjC family was one of five nodes that fit these criteria.

The H3K4me3 mark is linked to promoter activity and recruit transcriptional machinery^11–13^. KDM5B (JARID1B/PLU1) is a JmjC containing histone demethylase that removes H3K4 trimethyl marks to a monomethylated status and thus a global transcriptional repressor. The result of global repression is context dependent, and KDM5B has both tumor promoting^16,42,86^ and tumor suppressive activites^33,37,45^.

While both MYC and KDM5B determine cell fate, influence genome-wide programs, and are master regulators of hematologic malignancies, the coordination between these master regulators of gene expression is not well-characterized. KDM5B forms a complex with TFAP2C and MYC to repress p21, resulting in enhanced proliferation in MCF7 cells^87^. In drosophila, increased expression of the KDM5B ortholog Lid enhanced dMyc-dependent growth^88^, counter to our findings. Subsequent unreported findings reversed this conclusion, where dMyc actually suppresses Lid^89^, suggesting these TFs are discordant in function, consistent with our findings.

MYC is known to coordinate of the recruitment of HAT complexes such as p300 (H3K122ac, enhancing transcription), TRRAP associates with HAT complexes (NuA4). These interactions enhance transcriptional output. In contrast, KDM5B represses transcriptional targets. Further, KDM5B expression is not a result of MYC-induced transcriptional amplification, as its expression patters are inversely correlated.

In this study we demonstrate that MYC suppresses KDM5B. Our investigations do not show coordination or implicate heteromeric formation between MYC and KDM5B. Instead, we observe that MYC suppresses KDM5B across several lymphoma and leukemia models. When MYC expression is revoked or inhibited across multiple murine and human hematopoietic malignancies, we detected restoration of KDM5B levels (Fig. 2B, 3D). In a blinded study of DLBC clinical samples we find that high MYC expressing cells have little to low KDM5B signal, and the inverse expression patterns are also observed (Fig. 3E)

Further, we note that while MYC binding site recognition is inclusive of the E-Box (CACGTG)^90^, the primary determinant for MYC binding sites is chromatin recognition with the H3K4me3 mark having the strongest association^91^. KDM5B specifically demethylates H3K4me3 and K3K4me2 chromatin marks^92,93^. Thus, KDM5B governs chromatin marks that orient MYC-dependent directives and thus regulates the MYC network (Fig. 5B, 5C). Additionally, we also found that KDM5B directly regulates the H3K4me3 marks in the promoter of MYC. Taken together, KDM5B regulates not only the histone marks that direct MYC binding, but the MYC locus itself.

The overexpression of MYC in MEFs under growth limiting conditions has been shown to induce apoptosis, indicating that a cell intrinsic apoptotic feedback loop exists. It is likely that although this feedback loop is suppressed in cancer it remains intact for reactivation. This mechanism has proven to be elusive. In MYC-dependent models of cancer, withdrawal of MYC results in apoptosis, likely due to this cell intrinsic feedback loop. In KDM5B KO cells the withdrawal of MYC shows little to no increase in apoptosis (Fig. 4B), and a more aggressive phenotype in vivo (Fig. 4D). Thus, the MYC/KDM5B axis contains a negative feedback loop that controls apoptotic responses.

We submit that KDM5B expression serves as a barrier to MYC-dependent malignant transformation, and is therefore suppressed in our cancer models. The expression of MYC is tightly controlled in normal cells with half-lives of about 30 minutes, although the restoration of KDM5B expression is much delayed considering this timeframe. We also did not observe direct MYC binding to the KDM5B locus (Fig. 6C), nor does the MYC/Miz1 suppressive complex influence KDM5B expression (Fig. 2F). Our data supports an indirect, albeit MYC dependent, regulation of KDM5B. Future studies to find this target or complex raise interesting therapeutic strategies to restore KDM5B expression and reinstate the intact negative feedback loop between MYC and KDM5B (Fig. 6D).

Aside from finding the unknown intermediary of KDM5B suppression several additional questions remain. Is the KDM5B enzymatic function required for suppression of the MYC network or is KDM5B a member of a larger complex that serves as a rheostat on MYC expression? Is the transcriptional amplification attributed to MYC invasion of the transcriptome a permitted through a coordination recruitment of HATs to open chromatin, and the suppression of the KDM5B-mediate transcriptional repression? Are specific members of the JmjC family of histone demethylases implicated in our initial computational approach across multiple MYC-driven malignancies undergoing similar repression in specific cancer subtypes? Is the suppression of KDM5B necessary or sufficient for malignant transformation?

In summary, we utilized an inference computational model to discover critical programs that support multiple MYC-dependent malignancies. We find that the histone demethylase KDM5B is repressed in MYC-driven hematopoietic cancers. We demonstrate that KDM5B regulates MYC-dependent programs, regulates pro-apoptosis responses through a negative feedback loop, and directly regulates the MYC locus.

## Declarations

### Ethics approval and Consent to Participate

Studies did not involve human subjects. Special consideration was given to minimization of pain and distress for the humane and ethical treatment of the animals utilized for the study. In addition to coordinating with APLAC protocols and veterinarians and obtaining IACUC approval, each experiment was considered for whether the number of mice utilized is appropriate, and consideration of humane endpoints to reduce pain and distress.

### Availability of data and materials

The datasets used and analyzed during the current study are available on the central repository GEO: GSE 51011, GSE 23743, GSE 36354, GSE 44672. The KDM5B KO RNA-seq and ChiP-seq data will be deposited upon publication of the manuscript.

## METHODS

### Cell culture conditions

Lymphoid cells lines were cultured in RPMI with L-glutamine (Gibco) supplemented with 10% fetal bovine serum (Tissue Culture Biologicals) with penicillin/streptomycin (Gibco) in a humidified 5% CO_2_ atmosphere. Cells were maintained at 37°C in a humidified incubator with 5% CO2, and typically passaged every three days.

### RNA extraction & cDNA synthesis

RNA extraction from 2 × 10^7^ cells is done using the Qiagen RNEasy Extraction kit or a Zymo Research Quick-RNA Miniprep kits. RNA quality and concentration are assessed by a spectrophotometer, the Nanodrop. cDNA is then synthesized from 0.4 μg of the extracted RNA using Qiagen cDNA reverse transcription kit. The cDNA is then stored at −20°C.

### qPCR

Primers are designed by using NCBI PrimerBlast program and primer specificity was then verified using BLAST. Primers were generated by the Stanford PAN facility. Real-time PCR is performed in 96-well plates on an ABI Biosystems Thermo Cycler 7500. All primers are detected by using SYBR Green as fluorophore. Reactions are carried out in 20 μl that contained 1.5 μl cDNA, 0.5 μM forward and reverse primers and 8uL water and 10 μl of 2× SYBR Green master mi (ABI). Amplification cycle is as follows: 95°C for 3 min, 35 cycles of 95°C for 10 s, 63°C for 30 s, 72°C for 30 s and a final extension at 72°C for 5 min. At the end of the amplification cycles, a dissociation curve is done to verify non-specific amplification. The thermal cycler software generated threshold cycle (Ct) values for each gene; Ct is the number of cycles required to reach the threshold fluorescence 15 standard deviations above the noise. The Ct values are exported into Excel for analysis, and GraphPad Prism for statistical analysis

### RNA-seq

MYC-dependent cells derived from the primary *Eµ-tTA/Tet-O-MYC* murine model and maintained in culture. 100 X 10^6^ cells were seeded in T175 flasks in fresh media and timepoint samples began 24hr post seeding. Cells were dosed with 20ng/ml doxycycline and samples collected at the indicated time-points. RNA extraction included QIA shredder step, RNEasy kit, and on column DNA digestion (all reagents from Qiagen). Samples were shipped on dry ice to Beijing Genomics Institute (BGI) for the RNA-seq QC and sequencing pipeline and read analysis.

### ChIP-Seq experiments

Libraries were sequenced on the NovaSeq platform (single-end, 100bp) at MPSSR (OHSU) and on the HiSeq 4000 platform at GC3F (UO). See table below for details. Each library was sequenced on multiple sequencing lanes, thus prior to alignment, reads from the same library were combined across lanes to obtain one raw read file per library. The quality and complexity of libraries were assessed following sequencing using the FastQC software (Andrews, 2010). Raw reads were aligned to the mouse reference genome (mm10) using Bowtie2 (Langmead et al., 2009) with default single-end settings. To investigate the overall success of ChIP-seq pull-down assays, we used the *mutliBamSummary* command, followed by *plotCorrelation* from the deeptools package (Ramirez et al., 2014) to calculate and visualize Pearson correlation coefficient between ChIP-seq datasets and hierarchical clustering of libraries based on similarity of the ChIP-seq signal genome-wide. Successful ChIP-seq assays are expected to show overall high correlation within the same histone mark.

#### Peak Calling and Fold-enrichment Calculations

MACS2 (Zhang et al., 2008) was used to identify significant (q<0.01) peaks against the corresponding input, in each dataset. Since the H3K27me3 histone mark form broad peaks in ChIP-seq experiments, libraries from these two marks were analyzed using the built-in *–broad* peak-calling setting in MACS2.

#### Differential histone Site Analysis

The deeptools program (Ramirez et al., 2014) was used to visualize overall differences in intensity (Reads Per Kb Million or RPKM) of histone marks at the TSS (+/−3Kb) of all genes in each of all four cell lines (Figure 3). This approach will allow visualizing gross genome-wide differences in the intensity of epigenetic marks, but may not allow detection of local differences occurring on a small subset of genes. To compare genome-wide epigenetic differences between cell lines, we used the diffReps program (Shen et al., 2013) to perform pair-wise comparisons of histone ChIP-seq assays. The diffReps program works by using a sliding-window approach to scan ChIP-seq signal genome-wide and compare it between “treatment” and “control” files. We used the default sliding window settings (window size of 1Kb and step size of 100bp) and statistical test (negative binomial) for ChIP-seq datasets. After the detections are done, diffReps merges the significant overlapping windows to define a the final “differential site” and re-evaluates the statistical significance. After significant differential sites for each of the pair-wise comparisons were identified, we used the BEDOPS program (Neph et al., 2012) to annotate the nearest gene to each differential site.

### CellTiter-Glo Assay

Cells were seeded at 10,000 cells per well in 96 well format in standard cell culture conditions described. 24 and 48 hours post seeding cells were treated as indicated. At 72 hours post seeding cells were assayed using CellTiter-Glo assay (Promega G7570). After a 5 min incubation, cell populations were measured using spectramax (Molecular Devices). Background values were subtracted (media + CellTiter-Glo) and average of 4 wells were normalized to vehicle treated cells. Statistical significance was determined using the Student’s T-test.

### CellTiter-Glo 2.0 Assay

Drug treatment: Cell proliferation was obtained using the Cytation5 (BioTek) as a luminescence reader. Cells were plated at 3,000 cells per well in 100µL of media, and then let grow for 24 hours in a 96 well white walled plate (Coring Incorporated Costar, 3610). After 24 hours cells were treated with TOFA (Cay10005263-1000, Cayman Chemical Company), Firsocostat (S8893, Selleck Chemicals), TBV-2640 (S9714, Selleck Chemicals), or CP-640186 (PZS0362, Millipore Sigma) at the following concentrations: .1, .3, 1, 3, 10, 30 µg/ml. After either 24 hours or 48 hours of growth under treatment 50µL of CellTiter-Glo 2.0 (G9242, Promega) is added to each well to be read. The plate was then rocked on a shaker for 2 minutes and then taken off the shaker and incubated at room temperature for 10 more minutes. The plate was then read on the Cytation5 (BioTek) at 6 luminescence fibers at 135nm for 1 second per well.

### Annexin V and 7-AAD experiments

Cells were seeded in T25 flasks and treated as indicated. At 24hrs post seeding/post initial treatment, remaining cells were treated to obtain 24hr timepoint. At 48hrs post seeding/post initial treatment cells were harvested for analysis. Briefly, 1 million cell aliquots from each treatment were collected and washed twice in cold PBS, followed by centrifugation at 1300 RPM. Cells were resuspended in 1X Annexin V Binding Buffer (BD Pharmingen). Cells were then stained with either no stain, 7-AAD (BD cat# 51-68981E), or PE Annexin V (BD cat# 5165875X) or 7-AAD and Annexin V for one hour. Cells were then detected by flow cytometry (using BD FACS Aria SORP instrumentation). Data Analysis was performed using FlowJo (TreeStar).

### *In Vivo* Mouse Experiments

Cell lines generated from *Eµ-tTA/Tet-O-MYC* transgenic murine model of MYC dependency in T-ALL counted and 2 million cells subcutaneously injected into NSG mice. Tumors were measured every two days until tumor burden reached terminal endpoint of 2 cm, whereupon mice were then dosed with doxycycline and further tracked for tumor relapse.

### Oncogene inactivation

Conditional MYC-driven mouse T-ALL cell lines were derived from Em-tTA/tet-O-MYC mice1. c-MYC was inhibited in *Eµ-tTA/tet-O-MYC* T-ALL and P493-6 cells by treating cell cultures with 0.02μg/ml doxycycline (Sigma-Aldrich, T7660) for indicated time-points. The conditional cell lines were confirmed to be negative for mycoplasma contamination and maintained in Roswell Park Memorial Institute 1640 medium (RPMI, Invitrogen) with GlutaMAX containing 10% fetal bovine serum, 100 IU/ml penicillin, 100 μg/ml streptomycin, 50 mM 2-mercaptoethanol at 37°C in a humidified incubator with 5% CO_2_.

